# Free long chain fatty acid solitarily primes early postembryonic development in *Caenorhabditis elegans* under starvation

**DOI:** 10.1101/2023.03.14.532521

**Authors:** Meiyu Ruan, Fan Xu, Na Li, Fukang Teng, Huanhu Zhu

## Abstract

Postembryonic development of animals is long considered an internal predetermined program, while macronutrient is essential only because they provide biomatters and energy to support this process. However, in this study, by using a nematode *Caenorhabditis elegans* model, we surprisingly found that dietary supplementation of palmitic acid alone, but not other essential nutrients of abundance such as glucose or amino acid mixture, sufficiently initiated the early postembryonic development under complete macronutrient deprivation. Such a development was indicated by changes in morphology, cellular markers in multiple tissues, behaviors and the global transcription pattern. Mechanistically, palmitate doesn’t function as a biomatter/energy provider, but as a ligand to activate the nuclear hormone receptor NHR-49/80 and generate an obscure peroxisome-derived secretive hormone in the intestine. Such a hormonal signal was received by chemosensory neurons in the head in regulating the insulin-like neuropeptide secretion and its downstream nuclear receptor to orchestrate the global development. Moreover, the nutrient-sensing hub mTORC1 played a negative role in this process. In conclusion, our data indicate that free fatty acid acts as a prime nutrient signal to launch the early development in *C. elegans;* and implicate that specific nutrient rather than the internal genetic program is the first impetus of postembryonic development.

## Introduction

The initiation factors of embryonic development seem very well studied (such as fertilization or parthenogenesis). However, whether there is a counterpart in postembryonic development was not clear. Nutrients are no doubt important environmental factors in this process. Under favorable nutrient conditions, animals grow at the optimal speed (Rashid et al., 2020). When experiencing the nutrient deficiency, the early postembryonic developmental process is usually slowed, or even arrested (diapause) until the nutrient condition is improved(Artyukhin et al., 2013). In the last decades, scientists have discovered multiple nutrient-sensing signaling pathways, such as the insulin and insulin-like growth factor signaling (IIS), AMPK and mTOR pathways, that critically regulate cell/animal growth during postembryonic development(Chantranupong et al., 2015; Gonzalez and Hall, 2017; Kim and Guan, 2019). It seems to be a common view that the cell/animal growth starts only if these major nutrient-sensing pathways are jointly activated by multiple macronutrients (such as carbohydrates, amino acids, or lipids) and even some essential micronutrients (Qi et al., 2017). In other words, the availability of each essential macronutrient acts as an AND-gate to initiate the development(Valvezan and Manning, 2019). Theoretically, such an AND-Gate model is logical and should be conserved in vivo because cells or animals that grow without any major essential macronutrient will die eventually. However, some reports indicated an alternative situation in vivo. For example, rats grow without essential unsaturated fatty acids started to show skin defects in 3-4 weeks (indicating the nutrient deficiency), but were still able to grow, gain weight and survive for at least 5-6 months before dying of kidney failure(Burr and Burr, 1929; Burr and Burr, 1930). Therefore, it seems that only part of key macronutrients contributes to the developmental decision in animals. However, experimentally proving such a hypothesis in vivo by withdrawing specific macronutrient (we called it a deductive strategy(Zhu et al., 2021b)) would be less convincing, because it could not distinguish their role as a regulator, or simply an essential biomatter/energy supplier in the development. In addition, it was also technically impractical since most animals will quickly die when a certain essential nutrient was missing in the food.

Nematode *C. elegans* is an ideal model organism to study such a question. First, like many other organisms, the optimal development of *C. elegans* requires many nutrients, such as glucose, amino acid, lipids, and several B vitamins (Lu and Goetsch, 1993; MacNeil et al., 2013; Qi et al., 2017; Zečić et al., 2019). Second and most importantly, *C. elegans* would enter a reversible developmental arrest named diapause and survive for a long period of time under nutrient insufficiency, or even complete macronutrient deprivation (we called it starvation hereafter for convenience)(Baugh, 2013; Baugh and Sternberg, 2006). Therefore, we can study the role of individual nutrients in *C. elegans* developmental regulation by simply dietarily supplementing them individually under the general nutrient insufficiency (we called it an inductive strategy(Zhu et al., 2021a)). By applying such a strategy, several previous studies including ours suggested that certain nutrients might act as a dominant signal to sufficiently promote the developmental process under the general nutrient insufficiency or starvation(Fukuyama et al., 2015; Qu et al., 2020; Zhu et al., 2021a). However, the full establishment of such a model needs more concrete evidence (See discussion).

In this study, we adopted and modified a previously reported L1 development assay to screen for a single nutrient that could initiate the early postembryonic development. By supplying individual macronutrient to newly hatched *C. elegans* under starvation, we surprisingly found only free long chain fatty acid such as palmitate, but not carbohydrates, amino acid mixture (AA mix), nor the combination of the latter two, initiated the very early postembryonic development. Our further mechanistic studies revealed that as a signal instead of a lipid metabolite, free long chain fatty acid orchestrates a sophisticated and cross-tissue regulatory program via unusual roles of the mTORC1, nuclear hormone receptors, peroxisomes, sensory neurons, and neuronal hormones in promoting postembryonic development.

## Results

### Palmitic acid specifically initiates *C. elegans* postembryonic development under starvation

We first established an improved assay to identify whether a single nutrient could initiate the early postembryonic development of *C. elegans* under fasting. We took the advantage of a commonly used method to synchronize newly hatched *C. elegans* first stage larvae (L1) by suspending them in the M9 buffer, a saline-like medium without any organic nutrients such as ethanol(Kniazeva et al., 2015; Qu et al., 2020). Under this condition, almost all animals were strictly arrested at the early L1 stage (Kniazeva et al., 2015; Qu et al., 2020). Then we supplemented individual nutrients into the M9 and observed the animal development after 48 hours (Fig. 1A). Interestingly, we found that only palmitate, but not glucose or amino acid mixture (AA mix hereafter), made animals significantly grow bigger (Fig. 1B, C). To further distinguish between a real developmental progression from a simple gain of body size, we checked a developmental marker, the maturation of AWC sensory neurons by observing the str-2::GFP expression pattern (Fig. S1A). In early L1 arrested animals, str-2::GFP showed a dim and symmetric pattern (AWC^off^), while in developing L1 animals, it became much brighter with an asymmetric, or occasionally symmetrically bright pattern (AWC^on^) (Fig. S1A)(Kniazeva et al., 2015; Troemel et al., 1999). We found only palmitate, but not the glucose, amino acid mix, nor even combining them together, could initiate the AWC maturation (Fig. 1D, E); Additional dietary supplementation of glucose, AA mix, or them together with palmitate also did not make a major difference in palmitate-induced AWC maturation (Fig. S1B). Moreover, a classical chemotaxis experiment (Bargmann et al., 1993a; Hsieh et al., 2014) also indicated that only palmitate-fed animals had maturated and functional AWC neurons to respond to the chemical cue butanone (Fig. 1F, S1C). In addition, to confirm that the palmitate-driven AWC neuronal maturation was the result of a real developmental event instead of a stress response that only affected neuronal tissues, we used another epithelial developmental marker *ajm-1::GFP*, which labels the division status of hypodermal cells (Fig. S1D, E) (Kaplan et al., 2015b; Knight et al., 2002). We found that palmitate, but not glucose or AA mix fed animals showed a significant hypodermal cell division (Fig. 1G, H). Interestingly, none of the palmitate-initiated early developmental events proceeded to about 6.3 hours after egg laying cultured under the normal NGM/*E. coli* condition (6.3h AEL), determined by seam cells, Q cells and M cells division markers (Fig. S1D-G) (Harfe et al., 1998; Kaplan et al., 2015b; Zheng et al., 2018), which suggest that the palmitic acid triggered a postembryonic development at the very early stage of L1 (Fig. 1I). Finally, our transcriptome analyses also confirmed that the gene expression pattern of palmitate-fed animals was largely different from fasting or glucose/amino acid mix supplemented animals, but shared a similarity with well-fed animals (Fig. 1J, S5A). Taken together, these data showed that free palmitate could single-handed initiate *C. elegans* postembryonic development (we named this event “<u>F</u>ree palmitate initiated <u>E</u>arly postembryonic <u>D</u>evelopment <u>U</u>nder <u>S</u>tarvation”, or FEDUS hereafter) in *C. elegans*.

**Figure 1.**
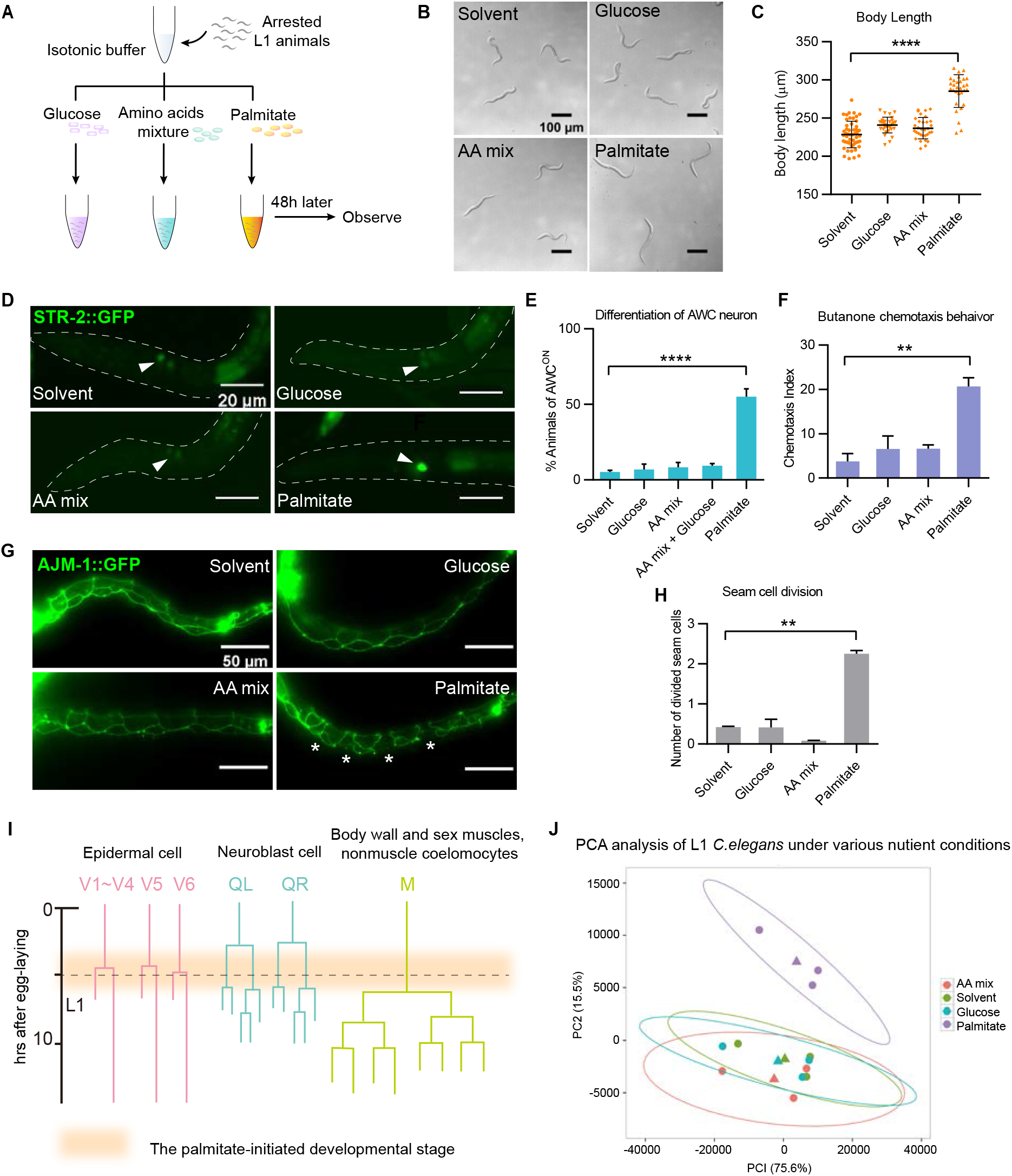
Palmitic acid promoted the development of arrested L1 *C. elegans* in the isotonic M9 buffer. (A) A schematic chart of our experimental design for screening key nutrient that initiates postembryonic development in *C. elegans*. Synchronized arrested L1 animals were cultured in M9 buffer with different nutrients and observed after 48-60 hrs. (B, C) Microscopic images (B) and a statistics graph (C) of the body length of the *C. elegans* cultured under various nutrient supplementation. (D, E) Fluorescent microscopic images (D) and a statistics graph (E) showing the percentage of animals with maturated AWC neurons cultured under various nutrient supplementation. AWC neurons (marked by *Pstr-2::gfp*) were indicated by arrowheads. Maturation of AWC neurons was indicated by intense and asymmetry expression *Pstr-2::gfp*. (F) A bar graph showing the chemotaxis behavior of *C. elegans* in response to butanone, an attractive odor detected by AWC neurons. (G, H) Fluorescent microscopic images (G) and a statistics graph (H) showing the development of seam cells, marked by AJM-1::GFP. Divided seam cells were indicated by asterisks. (I) A cartoon picture illustrating cell lineages of V1∼V6 seam cell, Q cell and M cell in the early postembryonic developmental stage. The orange color showed the time window of palmitate-induced development. (J) A chart showing PCA analysis of global gene expression among animals cultured under various nutrient supplementation. The palmitate-fed group (purple) was prominently different from the other three groups.

### mTORC1 negatively mediated palmitate-induced postembryonic development under fasting

We next investigated the responding tissues and related signaling pathways involved in FEDUS. First, we found anesthetics (such as the glutamate-gated chloride channel inhibitor ivermectin, or the nicotinic acetylcholine receptor agonist levamisole) that block food uptake of *C. elegans* by paralyzing the pharynx pumping, could significantly suppress the FEDUS (Fig. 2A, S2A, B). These data suggest that the palmitate needs to be ingested in the intestine, rather than directly sensed by sensory neurons to promote the postembryonic development. Previously, we have found that under the well-fed or dietary restriction condition, intestinal activation of mTORC1 (a central hub of nutrient-sensing machinery (Blackwell et al., 2019; Gonzalez and Hall, 2017; Saxton and Sabatini, 2017) was critical to promoting *C. elegans* L1 development (Kniazeva et al., 2015; Zhu et al., 2015; Zhu et al., 2013; Zhu et al., 2021a). Therefore, we tried to suppress the FEDUS by knocking down mTORC1. However, we found mTORC1 knocking down by *raga-1(-)* [the ortholog of mammalian RagA/B] or *daf-15(RNAi)* [the ortholog of mammalian Raptor] did not suppress the FEDUS (Fig. 2B, C), suggest that the activation of mTORC1 was not required for FEDUS. More surprisingly, we found *raga-1(-)* or *daf-15(RNAi)* was able to initiate the development even without the supplementation of free palmitate (Fig. 2B, C). We then measured the mTORC1 activity by the relative intestinal nucleolar size labeled by the FIB-1 antibody (because mammalian phosphorylated-S6K antibodies could not recognize its *C. elegans* ortholog RSKS-1) (Li et al., 2021; Ma et al., 2016; Sheaffer et al., 2008; Zhu et al., 2021a). Interestingly, we found the mTORC1 activity was further reduced, rather than increased under the palmitate supplementation (Fig. 2D, E). These data suggest that the inactivation of mTORC1 is sufficient to promote the development under starvation. Such an effect seems mTORC1 specific, because mTORC2 mutant rict-1(-) [orthologue of Rictor] did not cause a similar effect (Fig. S2C). To further confirm such striking findings, we constitutively activated mTORC1 by ubiquitously overexpressing RAGA-1^DA^, a dominant active form (Q63L mutation) of RAGA-1 (Mammalian RagA/B)(Zhu et al., 2013) and found it completely blocked the FEDUS (Fig. 2F). These data indicate that the inactivation of mTORC1 is necessary and sufficient to mediate the FEDUS.

**Figure 2.**
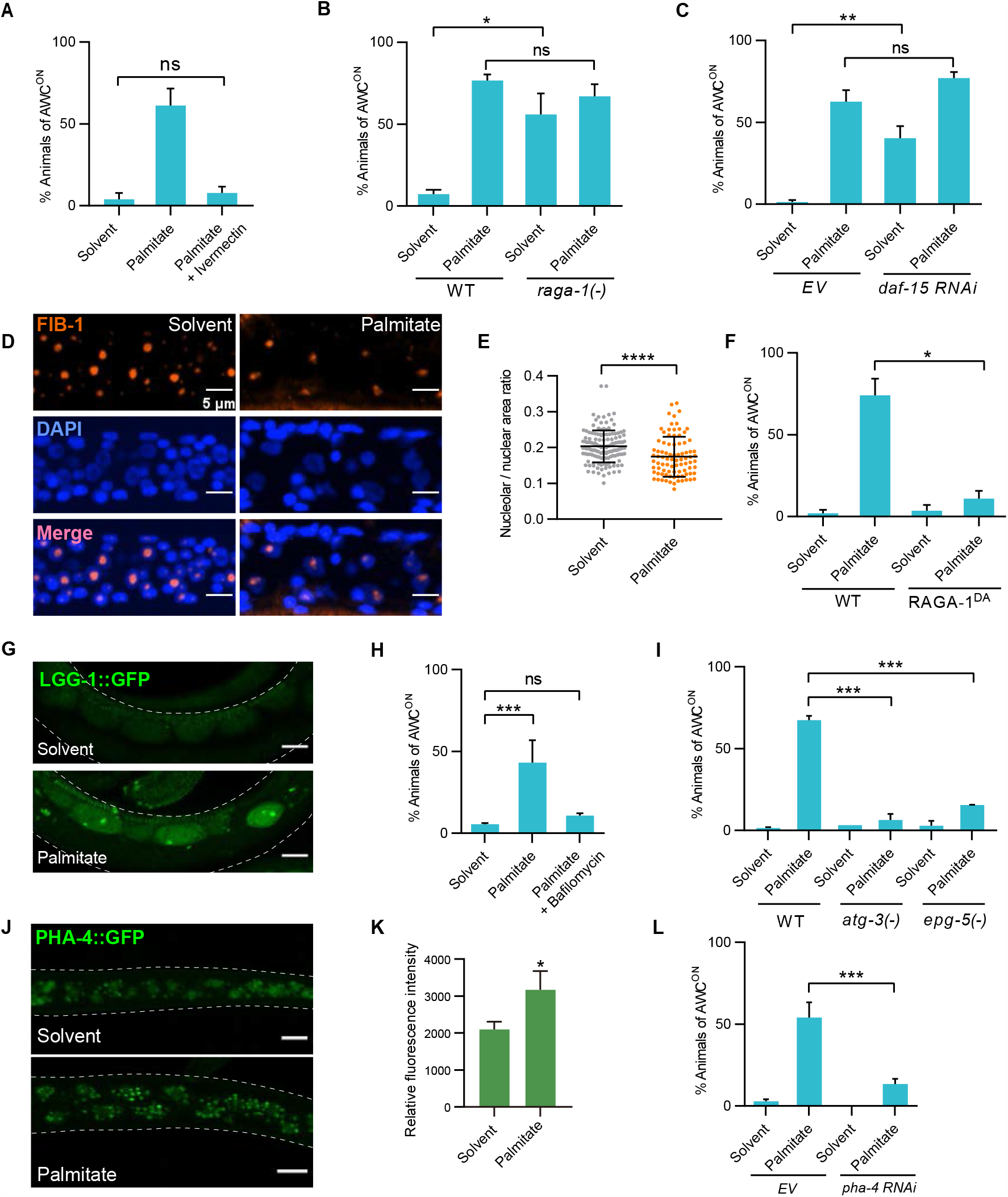
Inactivation of mTORC1 was necessary and sufficient to mediate the FEDUS. (A-C) (A) Anesthetics ivermectin significantly suppressed FEDUS. The loss function of *raga-1* (B) or RNAi of *daf-15* (C) facilitated AWC neuron maturation. (D, E) Representative immunofluorescent microscopic images (D) and statistical data (E) showing that the relative nucleolar size was significantly reduced in FEDUS. The nucleoli were labeled by the FIB-1 antibody (orange) and the nuclei were stained by DAPI (blue). (F) A bar graph showing a dominant-active mutant of RAGA-1^Q63L^ (RAGA-1 DA) suppressed AWC neuron maturation. (G) Fluorescent microscopic pictures showing the autophagy level indicated by LGG-1::GFP in epidermal cells. The autophagy was elevated in palmitate-fed animals (bottom). Dash lines marked the edge of *C. elegans* body. (H, I) A bar graph showing the percentage of animals with maturated AWC neurons under solvent or palmitate supplementation. Autophagy inhibitor bafilomycin (H), *atg-3*, or *epg-5* mutant (I) suppressed the AWC neuron maturation in FEDUS. (J, K) Fluorescent m. icroscopic images and statistical data showing the intestinal PHA-4::GFP expression. The level of PHA-4::GFP was increased in palmitate-fed animals. (L) A bar graph showing the percentage of animals with maturated AWC neurons under solvent or palmitate supplementation. RNAi of *pha-4* suppressed AWC neuron maturation in FEDUS.

### Autophagy is essential but not sufficient for palmitate/mTORC1(-)-induced postembryonic development

We then ask why the inactivation of mTORC1 by free palmitate triggered the development under starvation, given a common understanding that mTORC1 activity is relatively low under fasting (Kniazeva et al., 2015; Sheaffer et al., 2008; Zhu et al., 2015; Zhu et al., 2013; Zhu et al., 2021a). One plausible explanation is that the inactivation of mTORC1 triggered autophagy and generated amino acids and other nutrients from the lysosome-dependent recycling process upon starvation (Jung et al., 2010). Interestingly, a recent report suggests that, in mammalian tissue-cultured cells, the mTORC1 activity could be reactivated to a basal level under the prolonged starvation to attenuate autophagy, which preserves the capacity for long-term survival (Yu et al., 2010). However, the cause of such a reactivation was not clear(Yu et al., 2010). Therefore, we hypothesize that when the basal mTORC1 activity was inhibited by dietary-supplemented palmitate, the autophagy level could be restored and the autophagy-derived AA could be used to support the animal development. Indeed, we found the autophagy marker LGG-1::GFP was significantly upregulated (Fig. 2G). In addition, blocking autophagy chemically (by bafilomycin), or genetically (by mutation of *atg-3/ATG3 or epg-5/EPG5)*, also completely suppressed FEDUS (Fig. 2H, I, S2D). The upregulation of autophagy likely depended on PHA-4 (ortholog of FOXA), a transcription factor that is negatively regulated by mTORC1(Sheaffer et al., 2008), because palmitate supplementation enhanced the PHA-4::GFP expression, and *pha-4(RNAi)* dramatically suppressed FEDUS (Fig. 2J-L). On the other hand, mutation of *aak-2/AMPK*, another autophagy regulator, only moderately affected the FEDUS (Fig. S2E). Interestingly, chemically enhancing the autophagy by trehalose (Mizunoe et al., 2018; Seo et al., 2018) or polyamine(Eisenberg et al., 2009; Morselli et al., 2011) was not sufficient to activate the development under the starvation (Fig. S2F, G), which is consistent with our finding that AA mix along could not initiate the development under a similar condition (Fig. 1B-H). These data suggest palmitate-induced autophagy is necessary, but not sufficient for FEDUS. Therefore, in addition to its role to generate recycled AA as biomaterials by autophagy, palmitate may also play a more important role to initiate the development.

### Palmitate initiates the postembryonic development independent of its metabolites

We then investigated whether palmitate acts as a simple nutrient (to provide biomaterials and energy), or acts as a signal molecule to trigger the development. We first determined the minimal concentration of palmitate to initiate FEDUS and found as low as 0.05mM of palmitate was sufficient to initiate the animal development (Fig. 3A, B). In the contrary, 30mM glucose, or 60mM AA mix could not promote a similar development (Fig 3C, D). Moreover, dietary supplementation of palmitate beta-oxidation metabolites such as acetyl-CoA or its cytosol precursor citrate [which provides energy, or building blocks for the fatty acid/glucose/cholesterol de novo biosynthesis], did not trigger a similar development (Fig. 3E, F). Blocking cytosolic Acetyl-CoA biogenesis by ATP citric acid lyase inhibitor (hydroxycitric acid tripotassium hydrate), or by acetyl-CoA synthase inhibitor1 (CAS# 508186-14-9) also could not suppress FEDUS (Fig. 3E, F). These data indicate palmitate neither acted as an energy supplier nor as the acetyl-CoA donor in FEDUS. In addition, we also found short chain fatty acids (SCFA), which derive from the palmitate beta-oxidation, were not the real development-promoting effector (Fig. 3G), even under a much higher concentration (Fig. S3A). Similarly, ketone bodies (such as beta-hydroxybutyrate [βHB] or acetoacetate), a class of metabolites biosynthesized from free fatty acids and acting as a major energy source under the dietary restriction condition(Edwards et al., 2014), were also not the major reason of FEDUS (Fig. 3H, S3B). These data suggest that palmitate did not initiate the development mainly via its common catabolic metabolites under the fasting condition.

**Figure 3.**
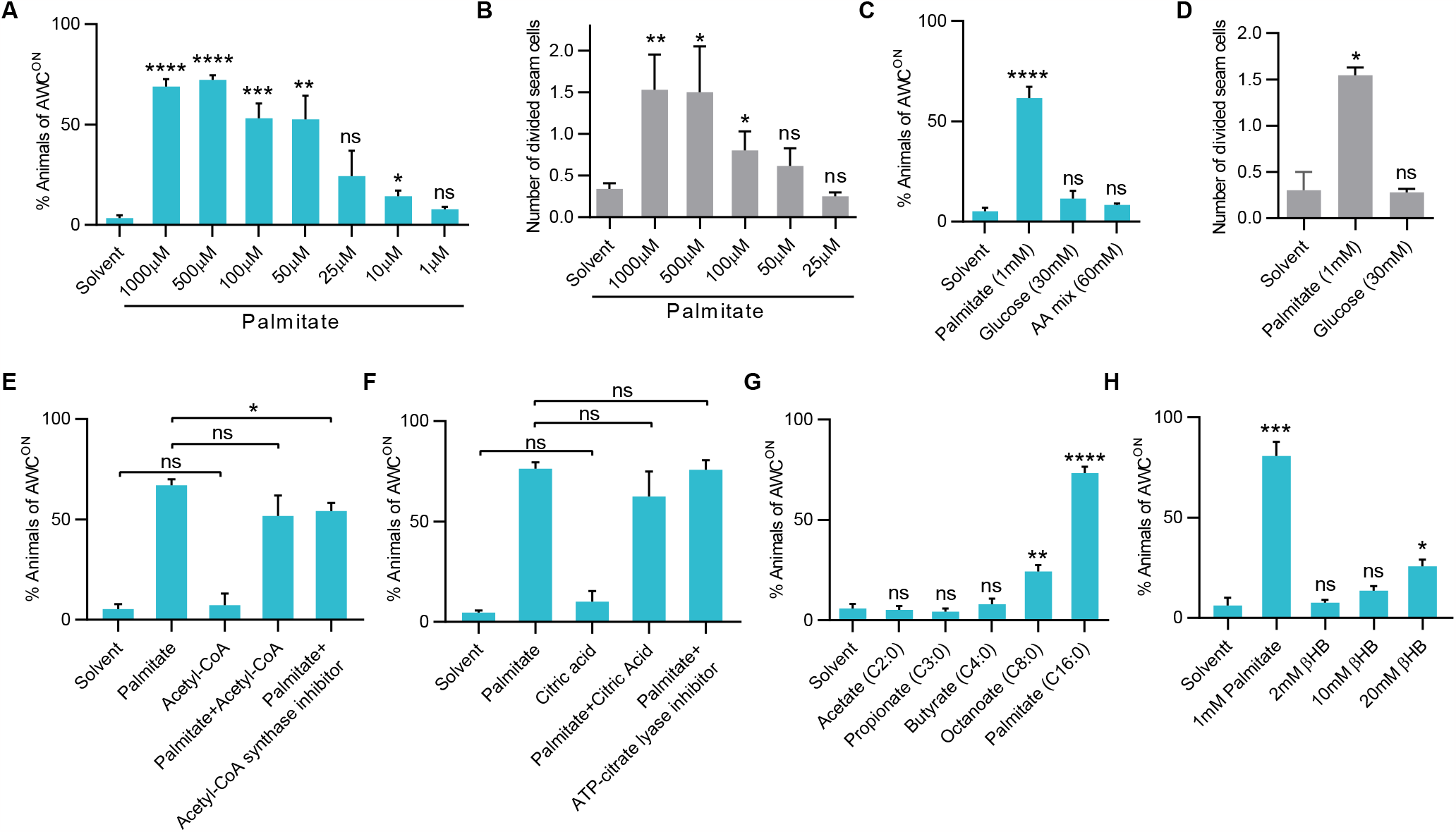
The development was not attributed to high metabolic energy. (A, B) Bar graphs showing the percentage of animals with maturated AWC neurons (A), or the average number of divided seam cells (B). Gradient concentration of palmitate promoted the maturation of AWC (A) and seam cell division (B). (C) A bar graph showing the percentage of animals with maturated AWC neurons. Dietary supplementation of 30mM glucose or 60mM amino acids mixture could not promote the maturation of AWC neurons. (D) A bar graph showing the average number of divided seam cells. Dietary supplementation of 30mM glucose was not able to promote the division of seam cells. (E-H) Bar graphs showing the percentage of animals with maturated AWC neurons under various metabolite supplementation. Neither Acetyl-CoA (E), citric acid (F), short chain fatty acids (G), nor the ketone body beta-hydroxybutyrate (βHB) significantly promoted AWC maturation. Neither Acetyl-CoA synthase inhibitor, nor cytosolic Acetyl-CoA ATP citric acid lyase inhibitor (hydroxy-citric acid) suppressed FEDUS.

Next, we tested whether the palmitate initiated the development via its anabolic metabolites. For example, many essential unsaturated fatty acids are derived from palmitate by a series of dehydrogenation(Vrablik and Watts, 2012; Watts and Browse, 2002; Zhu and Han, 2014). We found that genetically blocking long chain fatty acids dehydrogenases only showed moderate or no effect on FEDUS (Fig. S3C, D), suggesting that polyunsaturated fatty acid or their hormone metabolites (such as prostaglandin or endocannabinoids) were not required for FEDUS. Next, we excluded palmitate initiated the development via the biosynthesis of high order lipids such as glycerophospholipids and sphingolipids, or via protein lipidation (Kniazeva et al., 2012; Tang and Han, 2017); because we found genetic mutations of multiple acyl-CoA synthetases (key enzymes for the long chain fatty acid to be incorporated into high order lipid or protein lipidation) did not prominently inhibit the palmitate-induced development (Fig. S3E). These data suggest FEDUS was not mainly due to the anabolic metabolites of palmitate.

### Palmitate initiates the postembryonic development via nuclear-hormone-receptors-dependent peroxisome activation

The data above implicate palmitate itself, but not its metabolites, directly activate the development under fasting. Interestingly, we also found a variety of mono- or polyunsaturated fatty acids triggered the development under starvation (Fig. 4A), suggesting that long chain fatty acids (LCFAs) share a common function in the development initiation. LCFAs have been reported with multiple physiological roles in vitro and in vivo. We first excluded that LCFA acts as a uncoupler of the mitochondrial electron transport chain (Demine et al., 2019; Fedorenko et al., 2012) to initiate the development, because other mitochondrial uncouplers such as carbonyl cyanide m-chlorophenyl hydrazone (CCCP) could not induce a similar development like the FEDUS (Fig. S4A). Moreover, UCP1 inhibitor GTP, or the mutation of its *C. elegans* homolog *ucp-4*, could not inhibit the FEDUS (Fig. S4B, C). In addition, we also excluded LCFAs induced ROS(Baarine et al., 2012) as the main cause of FEDUS, because ROS inducer peroxide or ROS inhibitor N-acetyl-l-cysteine (NAC) couldn’t affect the FEDUS, and the expression of reactive oxygen species (ROS) or mitochondria unfolded protein response (UPR^mt^) marker HSP-6::GFP was not changed under FEDUS (Fig. S4D, E).

**Figure 4.**
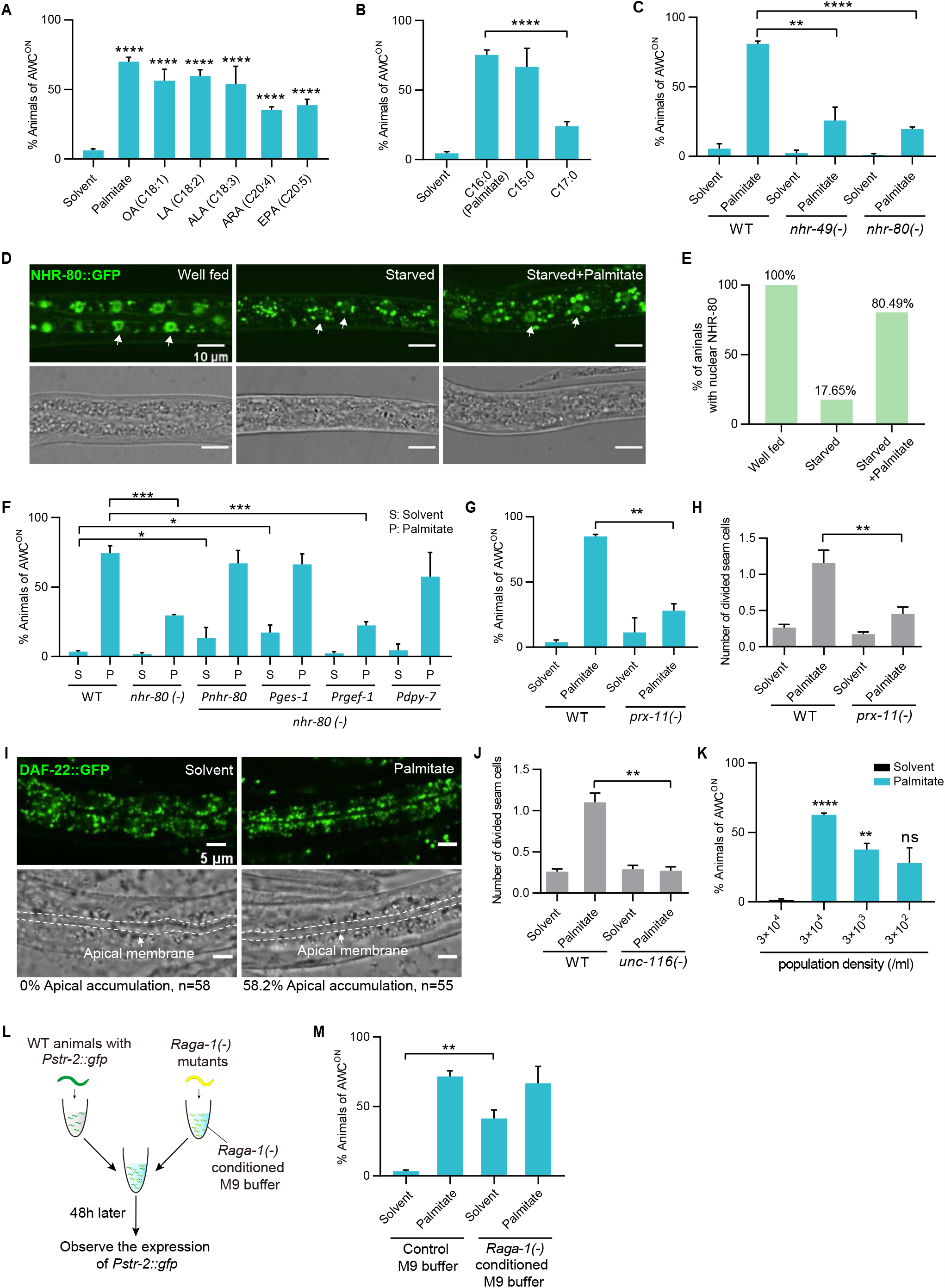
NHR-49/80 mediated the FEDUS via the peroxisomal activation. (A) Bar graphs showing the percentage of animals with maturated AWC neurons. (A) Dietary supplement of unsaturated long chain fatty acids such as oleic acid (OA, C18:1), linolenic acid (LA, C18:2), α-Linolenic acid (ALA, C18:3), arachidonic acid (ARA, C20:4) or eicosapentaenoic acid (EPA, C20:5) promoted the maturation of AWC neurons. (B) Dietary supplement of pentadecanoic acid (C15:0), but not heptadecanoic acid (C17:0), efficiently promoted the maturation of AWC neurons. (C) Loss function mutant of *nhr-49* or *nhr-80* significantly blocked the FEDUS. (D) Representative fluorescent microscopic images showing the subcellular localization of NHR-80::GFP under various nutrient conditions (upper panel). The nuclei were indicated by arrows. Related bright field microscopic pictures were also shown (bottom panel). (E) A bar graph showing the percentage of animals with nucleus-localized NHR-80::GFP. Palmitate supplementation dramatically increased the nuclear localization of NHR-80::GFP under starvation.. (F) A bar graphs showing the percentage of animals with maturated AWC neurons. Tissue specific expression of *nhr-80* in the intestine (*Pges-1*) or in the hypodermis (*Pdpy-7*), but not in neurons (*Prgef-1*) rescued the maturation of AWC neurons in *nhr-80(-)* animals. S is short for solvent. P is short for palmitate. (G, H) Bar graphs showing the percentage of animals with maturated AWC neurons (G), or the average number of divided seam cells (H). The loss function of *prx-11* inhibited both of them in FEDUS. (I) Representative fluorescent microscopic images showed the subcellular localization of peroxisomes marked by the *Pdaf-22::gfp::daf-22* knock-in strain. The same samples taken in the bright field were also shown. Dash lines highlighted the intestinal apical membrane. Palmitate supplementation significantly relocated peroxisomes to the intestinal apical region (the percentages of animals with apically localized peroxisomes were listed at the bottom) (J) A bar graph showing the average number of divided seam cells. The loss function of *unc-116* inhibited the seam cell division in FEDUS. (K) A bar graph showing AWC neurons maturation of animals grew at various population densities. Lower population density significantly decreased palmitate-induced AWC maturation. (L) A cartoon illustration of the experiment design that grow WT animals in M9, or *raga-1(-)* animals conditioned M9 (see method). (M) A bar graph showing that maturation of AWC neurons was significantly promoted in WT animals grew in *raga-1(-)* animals conditioned M9.

As the last resort, we tried two less common long chain fatty acids, pentadecanoic acid (C15:0), and heptadecanoic acid (C17:0), individually. We are surprised to find pentadecanoic acid, but not heptadecanoic acid efficiently initiated the postembryonic development (Fig. 4B), though the dietary supplemented heptadecanoic acid was efficiently absorbed and utilized by *C. elegans* (Fig. S4F). Such results immediately got our attention of the well-known nuclear hormone receptor PPARα since a recent paper reported that free fatty acid pentadecanoic acid, or palmitate, but not heptadecanoic acid, acts as agonists of mammalian PPARα in vitro and in vivo (Venn-Watson et al., 2020). Moreover, those unsaturated fatty acids (such as linoleic, linolenic acid, arachidonic, or eicosapentaenoic acid) that initiated FEDUS (Fig. 4A), are also known PPARα agonists (Krey et al., 1997). Furthermore, we found that inactive mTORC1 was essential for the FEDUS phenotype (Fig. 2B, C), which is consistent with a previous finding that mTORC1 inhibition was essential for PPARα activity (Sengupta et al., 2010). Last, we found genetic disruption of NHR-49, the *C. elegans* ortholog of PPARα, could significantly block the FEDUS (Fig. 4C). These data suggest PPARα/NHR-49 potentially mediates the FEDUS.

### PPARα/NHR-49 mediates the FEDUS via the peroxisomal activation

PPARα and its *C. elegans* ortholog NHR-49 are nuclear hormone receptors with pleiotropic functions such as regulating lipid metabolism, aging, and stress response (Antebi, 2015; Folick et al., 2015; Goh et al., 2018; Ratnappan et al., 2014; Van Gilst et al., 2005; Varga et al., 2011). NHR-49 usually needs other nuclear hormone receptors as the coactivator to assert its function (Pathare et al., 2012). We first determined NHR-80, the *C. elegans* homolog of mammalian hepatocyte nuclear factor 4 (HNR4), was also essential for FEDUS (Fig. 4C). Interestingly, N-acylethanolamines (NAEs), or deficiency of FAAH-1, an enzyme that degrading N-acylethanolamine (another ligand of NHR-49/80(Folick et al., 2015)), could not initiate FEDUS (Fig. S4G, H). These data suggest that the NHR-49/80-dependent FEDUS was specifically induced by free LCFAs.

We next elaborated where and how nuclear hormone receptors mediates the FEDUS phenotype. We found tissue-specifically restoration of NHR-80 in the intestine (driven by *ges-1 promotor*), or in hypodermis (by *dpy-7* promotor), but not in neurons (by *rgef-1* promoter), fully restored the FEDUS phenotype in *nhr-80(-)* animals (Fig. 4F). Furthermore, overexpression of NHR-80 ubiquitously or only in the intestine, was sufficient to partially initiated the AWC maturation in *nhr-80(-)* animals even without palmitate supplementation (Figure 4F). In addition, we found the nucleic localization of NHR-80 under the fed condition (Goudeau et al., 2011) shifted to the cytosol under starvation (Fig. 4D). Supplement of palmitate significantly increased the nuclear localization and the intensity of NHR-80::GFP (Figure 4D, E, S4I), Taking together, these data suggest that intestinal nuclear-localized NHR-80 critically mediates the FEDUS phenotype.

### FEDUS is mediated by a secretive hormone/perokine derived from apical intestine-positioned peroxisomes

Next, we investigated the major downstream pathway of NHR-49/80 to initiate the FEDUS, given several well-known functions of PPARα/NHR-49, such as gluconeogenesis, mitochondria beta-oxidation of free fatty acid, unsaturated fatty acid biosynthesis, and free fatty acid induced ROS detoxification (Antebi, 2006; Hu et al., 2018), have been excluded above (Fig. S4A-E). The remaining known function of PPARα/NHR-49 was to mediate peroxisomal proliferation, which is exactly what the name of PPAR comes from (Reddy et al., 1974). Interestingly, by analyzing our RNA-seq data, we found that peroxisome-related gene/pathways were highly enriched in the palmitate supplemented animals (Fig. S5B-E). Furthermore, blocking peroxisome function by mutation of *prx-11/PEX11*, which important for peroxisome proliferation(Li et al., 2021; Thieringer et al., 2003), also inhibited the FEDUS (Fig. 4G,H), even though *prx-11* was not essential for L1 development (Li et al., 2021; Thieringer et al., 2003; Wang et al., 2013). These data suggested that NHR-49/80 dependent peroxisome activation plays a critical role in FEDUS.

The critical role of peroxisome in FEDUS is very intriguing, especially because we recently have identified that *C. elegans* negatively regulated its L1 development via repositioning of peroxisomes to the apical region of intestine (e.g., close to the intestinal lumen) by Kinesin and facilitating a family of peroxisome-derived secretive metabolite hormones (Li et al., 2021). We then tested whether the FEDUS could also be regulated in a similar way. Surprisingly, we found that palmitate supplementation relocated peroxisomes to the apical intestine (Fig. 4I). Moreover, disruption of peroxisomal apical localization by a mutation of Kinesin heavy chain subunit UNC-116 completely disrupted the FEDUS phenotype (Fig. 4J)., suggesting apical localization of peroxisome is essential for FEDUS. Finally, we found the penetrance of FEDUS was positively regulated by the population density of the animals (Fig. 4K), suggesting that a secretive hormone plays an important role in FEDUS. To further verify such a possibility, we grew WT L1 animals with *raga-1(-)* animals (they could initiate their development without palmitate, see Fig. 2B) together in the isotonic M9 buffer (Fig.4L). We are surprised to find that even without palmitate supplementation, about 40% of WT animals started to develop (Fig. 4M); which strongly suggests a potential secretive hormone is sufficient to initiate the animal development under starvation. Taking together, these data indicate palmitate facilitates the secretion of a development-promoting hormone via relocating peroxisomes to the apical intestine and critically initiate the animal development.

We are very curious about the nature of such a peroxisome-derived hormone. PPARα/NHR-49 could activate the biosynthesis of multiple metabolites in peroxisomes, such as metabolites from the sterol de novo pathway, peroxisomal beta-oxidation or alkyl-glycerophospholipids. These metabolites could function as signals to mediate the FEDUS (Islinger et al., 2018; Li et al., 2021; Ludewig and Schroeder, 2013; Watterson et al., 2022; Xu et al., 2011). We first excluded metabolites in the de novo sterols pathway, because (1) supplement of several important intermediate metabolites in this pathway, such as mevalonate or coenzyme Q9 could not initiate the development under fasting (Fig. S4J, K); (2) inhibition the sterol de novo pathway by lovastatin that blocking the rate-limited enzyme HMG-CoA reductase, did not significantly affect the FEDUS (Fig. S4L). We also found peroxisomal beta-oxidation products, such as ascarosides were highly unlikely the signal, since blocking the peroxisomal beta-oxidation by a key thiolase mutant *daf-22(-)* also did not suppress the FEDUS (Fig. S4M). Finally, the fact that *fat-1(-)* failed to block the FEDUS phenotype ruled out the possibility that such a hormone was the recently identified small molecule hormone NACQ (Ludewig et al., 2019). Taking together, our data indicate that there is a type of new peroxisome-derived hormone, or “Perokine”, mediates the FEDUS phenotype.

### The Intestinal Perokine mediates FEDUS by targeting sensory neurons, downstream insulin-like neuropeptides and the DAF-12 pathway

Next, we studied how the development-promoting perokine further coordinated the whole-body development. *C. elegans* usually senses the hormone/pheromone via their ciliated sensory neurons (Inglis et al., 2007; Li et al., 2021; Li et al., 2022; Ludewig and Schroeder, 2013). We found disfunction of ciliated neurons by several mutations (*daf-6, xbx-1, or osm-6*) almost eliminated the FEDUS (Fig. 5A), suggesting this mystery intestinal peroxisomal signal is also sensed by ciliated sensory neurons. Because many neuropeptides are known to respond to nutrient availability to regulate the L1 development (Rashid et al., 2020), we analyzed whether FEDUS is dependent on the neuropeptide release to control the development. We found mutant of *unc-31(-)*, an orthologue of human CADPS (calcium-dependent secretion activator) which responsible for neuropeptide secretion (Lee and Ashrafi, 2008), almost completely suppressed the FEDUS (Fig. 5B, C, S6A). Moreover, *egl-3(-)* (human PCSK2) or *egl-21(-)* (human CPE), two enzymes that are essential for neuropeptide processing/production (Husson et al., 2006; Jacob and Kaplan, 2003), also blocked the FEDUS (Fig. 5D, S6A). These data suggest neuropeptides play a major role in FEDUS. Among many known neuropeptides, insulin-like signaling (IIS) are known to play essential roles in the early postembryonic development, especially under fasting (Baugh, 2013; Baugh and Sternberg, 2006). Thus, we disrupted IIS by using the IIS receptor daf-2 (ortholog of human insulin receptor) temperature-sensitive mutant *daf-2(e1370)* and found it completely inhibited the palmitate or *raga-1(-)* induced development under fasting (Fig. 5E). These data suggest that multiple insulin-like peptides synergistically act at the downstream of palmitate and mTORC1 to initiate the development.

**Figure 5.**
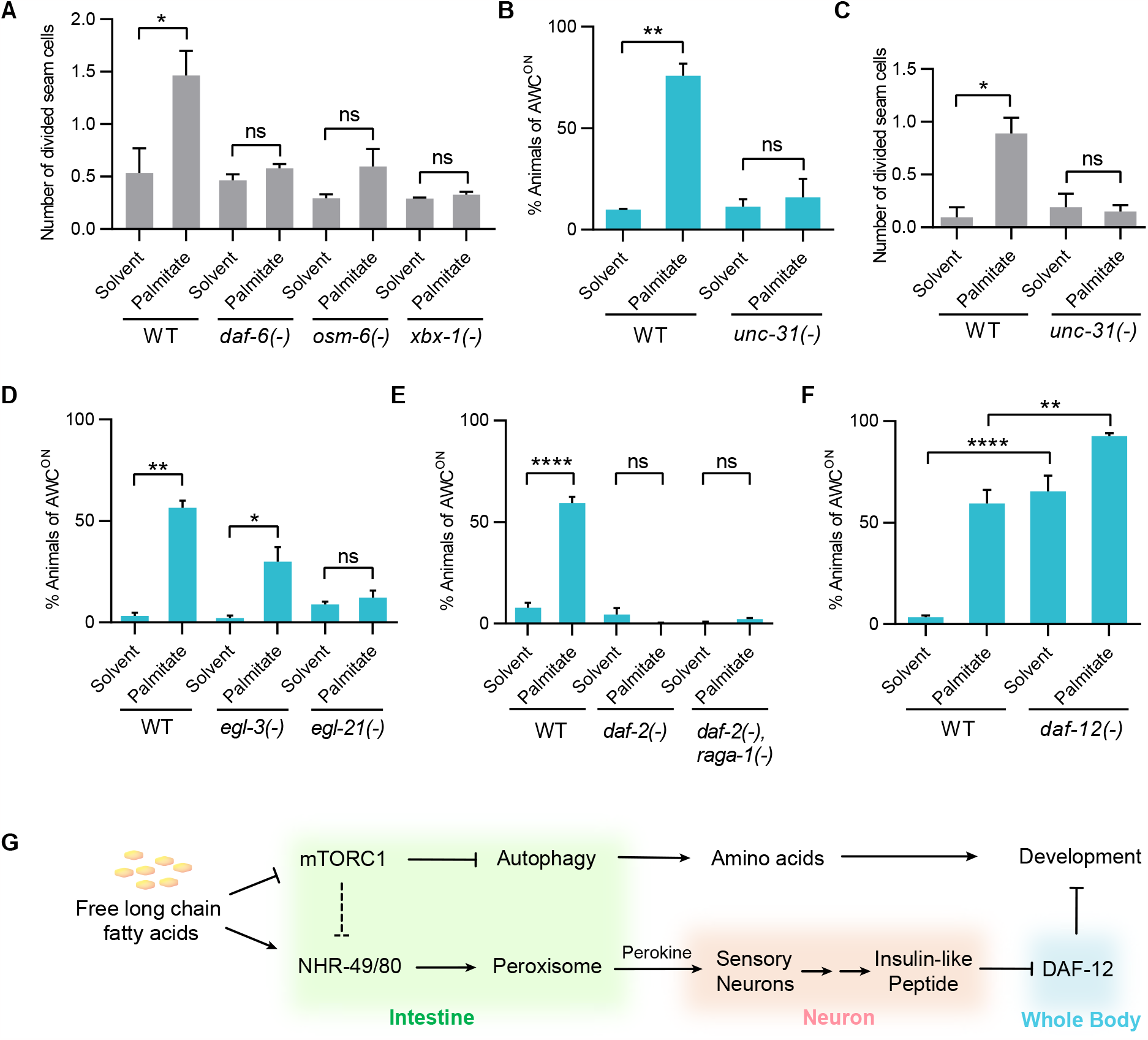
Ciliated sensory neurons and the insulin-like pathway mediated the FEDUS. (A) A bar graph average number of divided seam cells. The seam cell division was significantly decreased in mutants That block the function of ciliated sensory neurons such as *daf-6, xbx-1, or osm-6*. (B, C) Bar graphs showing the percentage of animals with maturated AWC neurons (B), or the average number of divided seam cells (C). The loss function of *unc-31* inhibited both of them in FEDUS. (D-F) Bar graphs showing the percentage of animals with maturated AWC neurons. (D) The palmitate-induced maturation of AWC neurons decreased significantly in neuropeptide processing/production mutants *egl-3* or *egl-21*. (E) A mutant of insulin-like receptor *daf-2* abolished the FEDUS of AWC neurons in control or *raga-1(-)* animals. (F) Loss function of *daf-12* promoted AWC neuron maturation with/out palmitate supplementation. (G) A cartoon model chart showing the cross-tissue mechanism of the free long-chain-fatty-acid-induced development under starvation (FEDUS).

The IIS pathway is known to execute its (developmental) function by transducing signal to transcription factor DAF-16 (mammalian FOXO) and nuclear hormone receptor DAF-12 (mammalian NR1I2) (Baugh, 2013). We further tested these two pathways and found *daf-12(-)* could initiate the L1 development under complete starvation (Fig. 5F). On the contrary, *daf-16(-)* was not able to initiate FEDUS by itself, a result consistent with the previous finding (Fukuyama et al., 2015). However, *daf-16(-)* was able to promote the development of fasting animals further, some even to the L2 larval stage (Fig. S6B). Therefore, these data indicate that the IIS-DAF-12 pathway critically mediated the FEDUS as the first postembryonic developmental checkpoint, while the DAF-16 pathway may play an important role in the next one. Taking together, we propose a model that free long chain fatty acids mainly act as a dominant nutrient signal to initiate *C. elegans* early postembryonic development via a sophisticated gut-brain axis (Fig. 5G).

## Discussion

Several previous works suggested that a few macronutrients could sufficiently activate the postembryonic development in *C. elegans*(Fukuyama et al., 2015) (Qu et al., 2020) (Zhu et al., 2021a), but following concerns made these studies far from conclusive. First, in some of those previous studies, multiple nutrients were needed to reactivate development, so one could not exclude the possibility of improved general nutritional status) (Fukuyama et al., 2015). Second, there were some nutrients still in the nutrient-insufficient or dietary restricted food so one cannot exclude the possibility that the supplementation of a specific nutrient molecule promotes the development just by increasing the efficiency of nutrient absorption/utilization(Zhu et al., 2021a). Third, the quantity of the supplemented nutrient was much higher than its physiological concentration, so the result may subject to alternative explanation(Qu et al., 2020). Last, the developmental assay was restricted to a specific type of cells, so that the cell division may result from a stress response rather than a physiologically-relevant developmental process (Qu et al., 2020),(Fukuyama et al., 2015). According, to overcomes these flaws by an optimized assay, we have discovered a surprisingly intriguing finding that LCFA such as palmitic acid, could initiate *C. elegans* postembryonic development without any other nutrients. One particular part of this finding is that other nutrient such as glucose or amino acid could not achieve a similar effect even under much higher concentrations (30mM glucose vs 50μm palmitate, 600X difference; 60mM total AA mix, 1200X difference) (Fig. 3C, D). Similarly, high concentrations of other nutrients/metabolites such as SCFA, citrate, or acetyl CoA also could not initiate the development (Fig. 3E-G). Such a huge difference could not be simply explained by an indirect effect from palmitate, such as a higher energy turnover efficiency, because free LCFA could be converted from other metabolites such as glucose or citrate endogenously. Interestingly, we found several triglyceride lipases, such as LIPL-3 and LIPL4, were also dramatically upregulated in palmitate-fed animals. These data not only suggest a feed-forward regulation of free LCFA in developmental regulation, they also imply that the upregulation of free long chain fatty acid level is a physiological process and possibly a rate-limiting step in the early postembryonic development. Interestingly, there is a significant amount of free LCFA in raw and commercially processed bovine milk (about 1-2 mM) and in milk fat (4mM) (Kintner and Day, 1965). It would be interesting to study whether those free LCFAs also play a critical role in the early postembryonic development in mammals.

Another interesting finding in this work is that the inactivation of mTORC1 is sufficient to initiate the FEDUS. In other words, mTORC1, a common developmental activator, plays a negative role in certain development stages. Very interestingly, Yu et al., have discovered that under prolonged starvation, cells across multiple species showed mTORC1 reactivation, possibly induced by autophagy-derived nutrients (Yu et al., 2010). They suggest such a pathway enables a negative feedback regulation of autophagy. However, the signal to initiate the autophagy-dependent mTORC1 reactivation and the physiological significance of such regulation remained elusive (Yu et al., 2010). Our work suggests that the depletion of free LCFA may trigger reactivation of mTORC1 and eventually arrested the animal under fasting. Therefore, animals would preserve their nutrient storage and enter the diapause stage, rather than quickly depleted their nutrient storage under starvation. We recently found that a specific type of LCFA, monomethyl-branched chain fatty acid, could activate the mTORC1 to restore *C. elegans* development from the L3 diapause under dietary restriction (Zhu et al., 2021a). It is very interesting to study whether free mmBCFA promotes the development via the inactivation of mTORC1 under complete starvation. We guess it could be due to the pleiotropic roles under different developmental stages (L3 vs early L1), or under different nutrient conditions (restricted food vs starvation). Illustrating the underlying mechanism would help us further understand the complicity of the nutrient-dependent developmental regulation in vivo.

Finally, our work proposed a new concept in animal development. The postembryonic developmental process in *C. elegans* seems not a pre-set program initiated mainly by its internal genetic machinery. Instead, just like mitosis, it proceeds with checkpoints gated by Postembryonic Developmental Control (PDC) genes, such as DAF-12, DAF-16 or LET-363/mTOR at various developmental stages respectively, and fundamentally initiated by specific Postembryonic Developmental Controlling Nutrient (PDCN), such as free LCFAs, amino acids or mmBCFA. If such a model is proven universally conserved among organisms, we may need a reevaluation of the concept of development and the role of nutrients. After all, the life system originated and evolved from those simple nutrients/metabolites.

## Acknowledgment

We thank Hong Zhang, Shohei Mitani, Gaofeng Fan, Yingchuan Qi, Yidong Shen, Shaobing Zhang, Hongyun Tang, Szecheng J. Lo, and Bin Liang for kindly supplied strains and suggestions. We thank *C. elegans* Knockout Consortium and National BioResource Project (NBRP) for mutant strains. We thank all of our laboratory members for many helps, especially Dr. Huiyang Xiong and Dr. Mengnan Zhu. We thank Ying Han, Piliang Hao from the Multi-Omics Core Facility (MOCF), Xiaoming Li, Ziwei Yang and Chengyu Fan from the Molecular Imaging Core Facility (MICF), Ying Xiong, Xiaoyue Ren from the Molecular and Cell Biology Core Facility (MCBCF) at the School of Life Science and Technology, ShanghaiTech University., Lei Zhang from Professional Technical Support Sharing Platform of Core facility (Improvement of Service Capability for Shanghai Proteomics Professional Technical Platform for Severe Diseases) of Shanghai Medical College at Fudan University for providing technical support. Funding: This work was supported by the National Natural Science Foundation of China (32170837)(HZ), the National Key R&D Program of China (2019YFA0802804[HZ], 2021YFA08004801[HZ]), the Recruitment Program of Global Experts of China (Youth)(HZ), Science & Technology Commission of the Shanghai Municipality (16PJ1407400)(HZ), Double First-Class Initiative Fund of ShanghaiTech University (SYLDX0162022)(HZ), and ShanghaiTech Startup program (HZ). Author contributions: MR, Conception and design, Acquisition of data, Analysis and interpretation of data, Drafting or revising the article; FX, FT, NL, Acquisition of data, Analysis and interpretation of data, Drafting or revising the article, Contributed unpublished essential data or reagents; HZ Supervised the study, Conception and design, Drafting or revising the article. Competing interests: Authors declare no competing interests. Data and materials availability: All data is available in the main text or the extended data materials.

## Materials and methods

### *Caenorhabditis elegans* strains and maintenance

*C. elegans* were maintained at standard techniques at 20°C (Brenner, 1974; Stiernagle, 2006) unless otherwise noted. N2 Bristol was used as the wild-type strain.

The following fluorescence strains were used: *kyIs140 [str-2::GFP + lin-15(+)], jcIs1(ajm-1::gfp), zdIs5 [mec-4::GFP + lin-15(+)], ayIs7(hlh-8::gfp), zcIs13(Phsp-6::hsp-6::gfp), DA2123[Plgg-1::gfp::lgg-1 + rol-6(su1006)], sydls031[Ppha-4::gfp-3*FLAG]*.

*The following mutant strains were used: raga-1(ok386), atg-3 (bp412), epg-5(tm3425), nhr49(nr2041), nhr80 (tm1011), prx-11(gk959960)*., *daf-22 (ok693), daf-6(e1377), osm-6(m201), xbx-1(ok279), unc-31(e928), egl-3(nr2090), egl-21(n476), daf-2(e1370), daf-12(rh61rh412), daf-16(mgDf50), aak-2(ok524)*,, *fat-1(ok2323), fat-2(wa17), fat-3(ok1126), fat-4(ok958), fat-5(tm420), fat-6(tm331), acs-22(tm3236), acs-19(tm4853), acs-20(tm3232), acs-17(ok1562), acs-2(ok2457), acs-7(tm6781), acs-14(ok3391), acs-5(ok2668), faah-1 (tm5011)*.

The following transgenic strains were used: *gfp-daf-22* knock-in strain was previously generated by our laboratory(Li et al., 2021). *Prpl-28::raga-1* vector was previously constructed by Dr. Zhu Mengnan in our lab(Zhu et al., 2021a). Plasmids were extracted by QIAGEN kit and injected to indicated *C. elegans* strains. Transgenic *C. elegans* were obtained by microinjecting corresponding plasmids (10ng/μl) into appropriate strains. 2ng/μl *pmyo-2::gfp* vector was used as the co-injection marker. Animals carrying the marker were considered transgenic strains.

### Macronutrient prepared

Fatty acids were all dissolved in DMSO and prepared as 10mM stocks. The solvent showed in most figures was DMSO except otherwise mentioned. Detail information of fatty acids was shown in Table S1.

The amino acid mixture and glucose were dissolved in ddH2O. The amino acid mixture was prepared according to the chemically defined *C. elegans* medium (CeMM) (Lu and Goetsch, 1993; Zhu et al., 2021a). The detailed recipe of the amino acid mixture is shown in Table S2.

### Microscopy

*Str-2::gfp, ajm-1::gfp, hlh-80::gfp, mec-4::gfp, hsp-6::gfp C. elegans* images were photographed by using the Olympus MVX-ZB10 fluorescence dissecting microscope and a BioHD-C20 CMOS camera. Autophagy analysis (*lgg-1::gfp*) was completed using Nikon CSU SORA spinning disk microscope. Images were acquired using a Nikon CSU SORA spinning disk microscope. *Pha-4::gfp* was photographed by Zeiss LSM 800 Confocal Laser Scanning Microscopy. Body length and GFP fluorescence were performed with Fiji software.

### Development assays

Eggs laid naturally by well-fed *str-2::gfp C. elegans* and L1 *C. elegans* on NGM plates were collected for growing to adult and subsequent bleaching. Noted that egg/L1 should be adjusted at a suitable number/plate to avoid food deprivation. Washes and bleaching were done according to standard protocols (Stiernagle, 2006). Synchronized L1 *C. elegans* in M9 buffer were divided evenly into different experimental groups. The fatty acid was dissolved in DMSO and prepared to 100mM stock solution. 3μl 100mM fatty acid solution was added into 200μl M9 buffer and immediately shake for 30-60s on vortex (experimental buffer). 100μl M9 with synchronized L1 *C. elegans* were added into the 200μl experimental buffer. The final concentration is 1mM. Noted that a high concentration of DMSO was harmful to *C. elegans* and needs to be used at a low concentration (No more than 1% volume ratio in this paper). 48h later, usually between 48h to 60h, the *str-2::gfp* expression was observed using a fluorescence dissecting microscope. Synchronized seam cell marker *ajm-1::gfp* L1 animals preparation was the same as the *str-2::gfp C. elegans*. The seam cell assay and statistical method were performed as previously described (Kaplan et al., 2015a). Above 50 animals were scored per replicate. At least three biological replicates were performed.

### Chemotaxis assays

Experiments were performed as described previously with modifications(Bargmann et al., 1993b; Park et al., 2005). 6cm plates prepared only with agar were used as chemotaxis assay plates. Above 100 treated L1 *C. elegans* were placed between the test spot and a control spot on the opposite side of the plate. 10 μl of 20 mM NaN3 was placed on both spots. After one hour, all plates were put into the 4°C refrigerator for the convenience of subsequent counting. *C. elegans* on the test/control spot were counted, and the chemotaxis index (C.I.) was calculated as the number of animals at the test spot minus the number of animals at the control spot, divided by the total number of animals (Fig S1B). A positive C.I. indicates attraction. At least three biological replicates were performed.

### RNAi experiment

The RNAi-feeding experiments were done as previously described (Kamath et al., 2001). dsRNA-expression constructs were from our previous works (Zhu et al., 2015) from the ORF-RNAi library (Open Biosystems) and sequences were verified. In all RNAi experiments, P0 *C. elegans* in the L1 stage (P0 L1 stage *C. elegans* were used in *elo-5* RNAi experiment) were seeded on the RNAi plates. After 3 days, adult *C. elegans* were bleached. F1 *C. elegans* were cultured in M9 buffer and assayed.

### RNAseq

Four animal groups with indicated dietary nutrient supplementation were performed the RNA-seq analysis: Solvent (DMSO), palmitic acid, glucose, amino acids mixture (AA mix). Three replicates were performed. Total RNA was prepared with TRIzol (Invitrogen). The sequencing and analysis were conducted by Biomarker Technologics Corporation (China).

### Immunofluorescence assay of FIB-1

L1 *C. elegans* were collected and washed three times with ddH2O. The immunofluorescence assay of FIB-1 was performed as previously described(Zhu et al., 2021a). The ‘‘frozen crack’’ method was used (Duerr, 2013). Fibrillarin (Abcam ab4566, 1:400) was used as the primary antibody, and goat anti-mouse conjugated to Alexa Fluor 594 (Invitrogen A11005, 1:800) was used as the secondary antibody. DAPI was used for staining nuclei. The intestinal cell nuclear (DAPI) and nucleolar (FIB-1) were quantified using Fiji software. The ratio of nucleolar/nuclear area was calculated.

### Lipid analysis by gas chromatography

Adult *C. elegans* cultured on above 100 plates (90mm) were collected and bleached. Bleached L1 were assayed in M9 buffer with solvent and C17:0 fatty acid each. After 48h, those L1 *C. elegans* in the M9 buffer were washed 3 times with ddH_2_O. After the last washing, water should be piped out as much as possible. The worm pellet can be frozen at -80°C. the total lipid extraction was based on a previous report (Bligh and Dyer, 1959). The method of gas chromatography was described in a previous report (Kniazeva et al., 2008).

### Statistical analysis

All statistical analyses except experiments, were performed using Student’s t-test using GraphPad Prism7 software. Data are presented as Mean ±SEM, and p<0.05 was considered a significant difference.

