## Supplemental figures for "Free long chain fatty acid solitarily primes early postembryonic development in *Caenorhabditis elegans* under starvation"

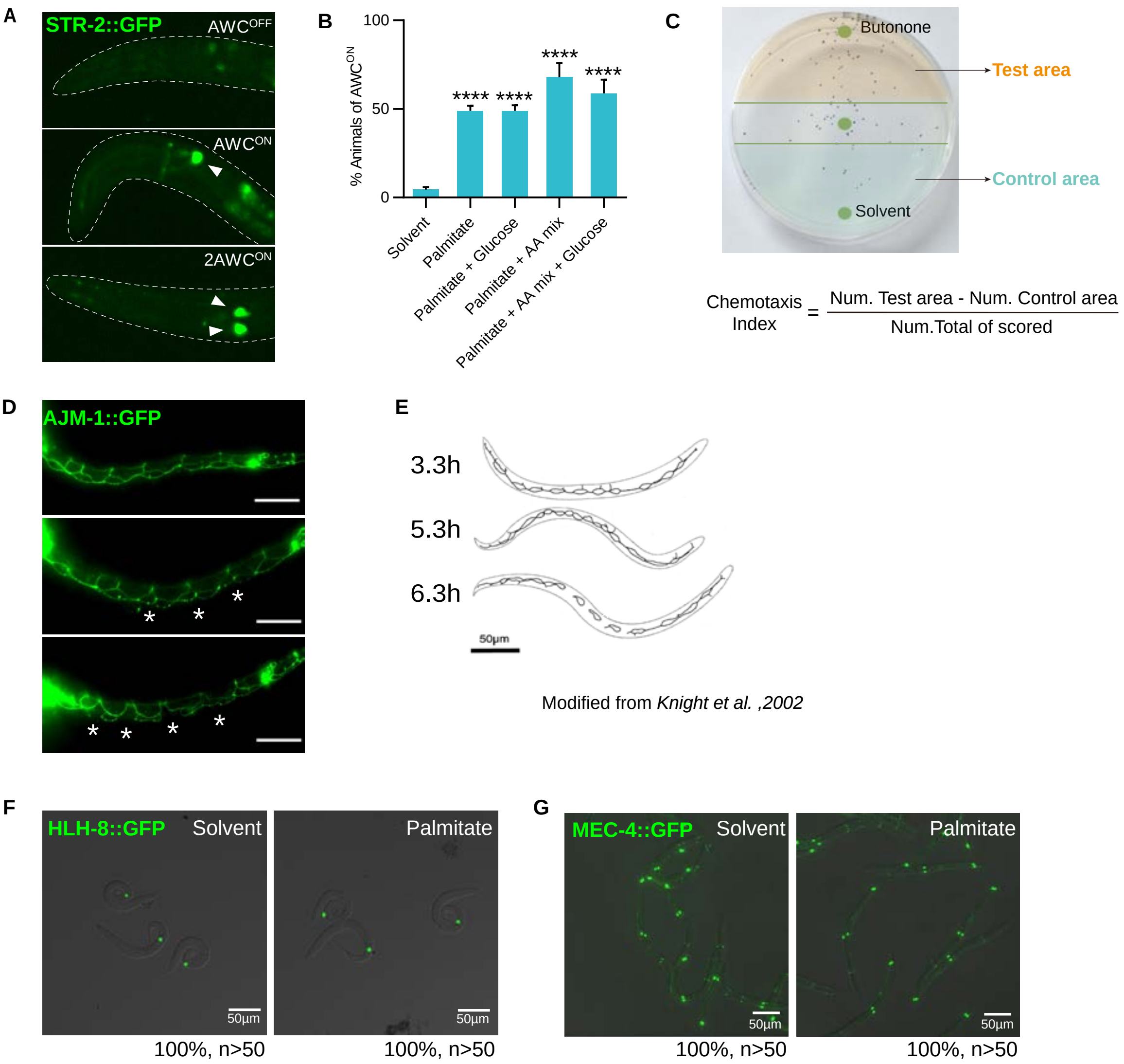


**Supplementary Figure 1.**
(A) Fluorescent microscopic pictures showing the maturation of AWC sensory neurons marked by *Pstr-2::gfp*. Maturated AWC neurons were indicated by arrowheads.
(B) A statistics bar graph showing the percentage of animals with maturated AWC neurons under various nutrient supplementation.
(C) A picture showing the chemotaxis assay to measure whether animals could be attracted to butanone on NGM plates.
(D) Fluorescent microscopic pictures showing the seam cells (marked by AJM-1::GFP) in palmitate-fed L1 animals. Divided seam cells were indicated by asterisks.
(E) A Cartoon picture modified from *Knight et al.* showing the seam cell development of *C. elegans* larvae at different time stages under the fed condition.
(F) Fluorescent microscopic pictures showing M cell division marked by HLH-8::GFP with/out palmitate supplementation. None of M cells was divided in both condition.
(G) Fluorescent microscopic pictures showing the Q cell division marked by MEC-4::GFP. No development difference was found between these groups.

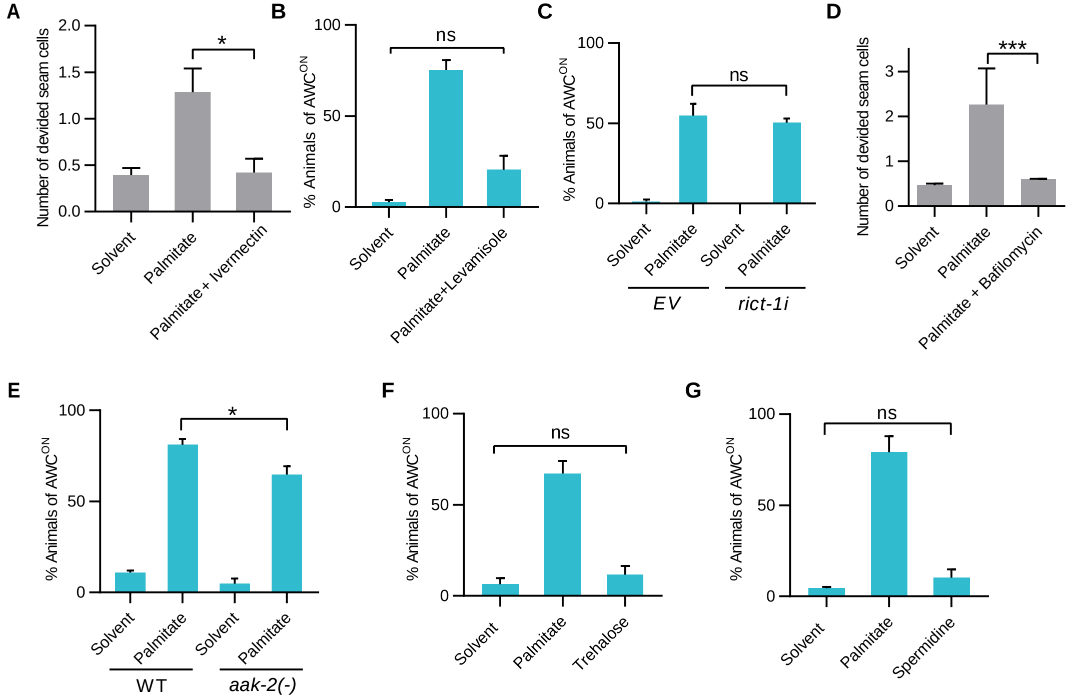


**Supplementary Figure 2.**

(A, B) A bar graph showing the average number of divided seam cells (A) and the percentage of animals with maturated AWC neurons (B). Animals treated with anesthetics, ivermectin(A) or levamisole (B), showed greatly decreased FEDUS.

(C) WT or *rict-1(-)* animal under solvent or palmitate supplement showed no difference in the AWC maturation.

(D) The autophagy inhibitor bafilomycin suppressed seam cell division in FEDUS.

(E) Loss function of *aak-2* could not suppress FEDUS, shown by the percentage of animals with maturated AWC neurons.

(F, G) Treatment of autophagy activator, 100mM trehalose (F) or spermidine (G) could not activate the AWC neuron maturation.

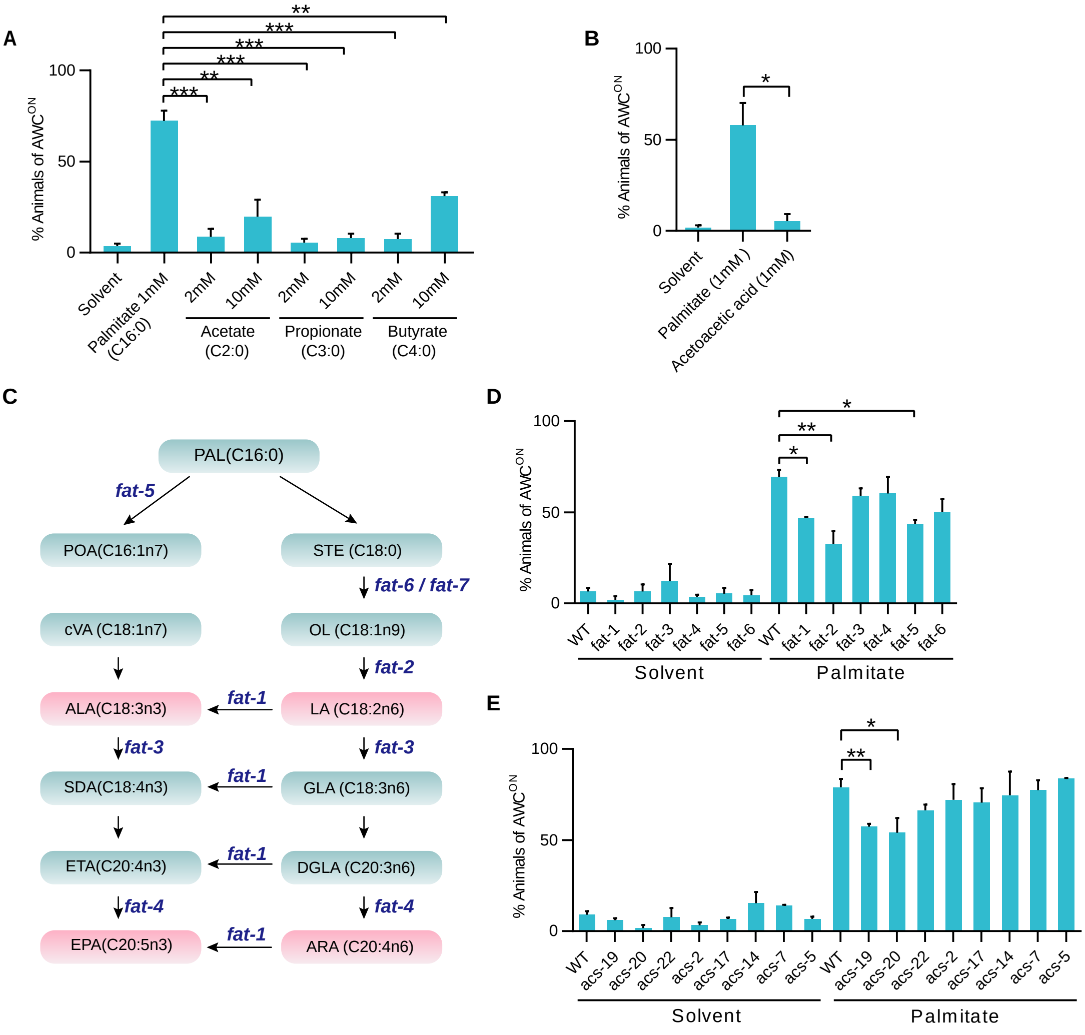


**Supplementary Figure 3.**

(A-B) Bar graphs showing the percentage of animals with maturated AWC neurons. Animals supplied with various concentrations of short chain fatty acids were tested (A). Supplementation of acetoacetate, a ketone body, could not initiate the AWC maturation (B).

(C) A cartoon illustration of the long chain fatty acid elongation pathway.

(D, E) Bar graphs showing the percentage of animals with maturated AWC neurons. (D) WT and multiple genetic mutants of long chain fatty acids dehydrogenases (FAT) in the PUFA biosynthetic pathway were tested. (E) WT and various mutants of Acyl-CoA synthase (ACS) under solvent or palmitate supplement were tested.

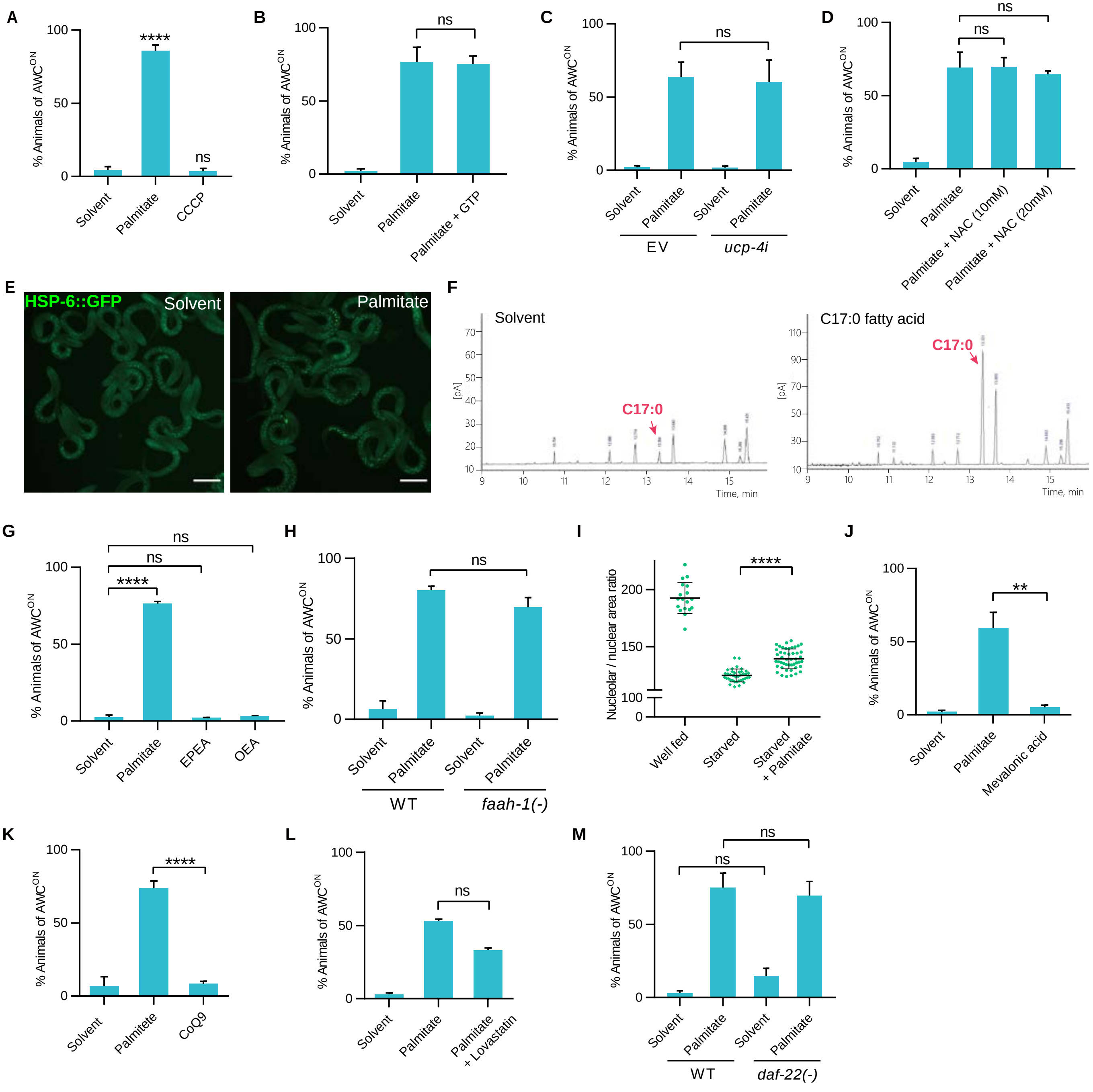


**Supplementary Figure 4**.

(A-D, G, H, J-M) Bar graphs showing the percentage of animals with maturated AWC neurons.

(A-D) A mitochondrial respiratory chain decoupler CCCP did not initiate the maturation of AWC neurons. Supplement of 1mM GTP(B) , the mutation of *ucp-4* (C), or the ROS inhibitor NAC (D) could not inhibit the maturation of AWC neurons.

(E) Representative fluorescent microscopic images showing the ROS level indicated by HSP-6::GFP under solvent or palmitate supplementation. There was no significant difference between these two groups.

(F) Gas chromatography picture showing the fatty acids profile of L1 animals. Dietary supplementation of heptadecanoic acid (C17:0, red arrows) was indeed absorbed by *C. elegans.*

(G) Supplement of EPEA (eicosapentaenoyl ethanolamide) or OEA (oleylethanolamide), two different NAEs, had no effect on the maturation of AWC neurons.

(H) Mutation of *faah-1* had no function on the maturation of AWC neurons.

(I) Statistical data showing that the relative fluorescence intensity of NHR-80::GFP was significantly downregulated in starved animals, while palmitate supplementation partially restored its expression.

(J, K) Supplement of mevalonic acid (J) or coenzyme Q9 (K) did not initiate the maturation of AWC neurons.

(L,M) Lovastatin, the HMG-CoA reductase inhibitor, or (M) a *daf-22* loss-of-function mutant, could not suppress the FEDUS.

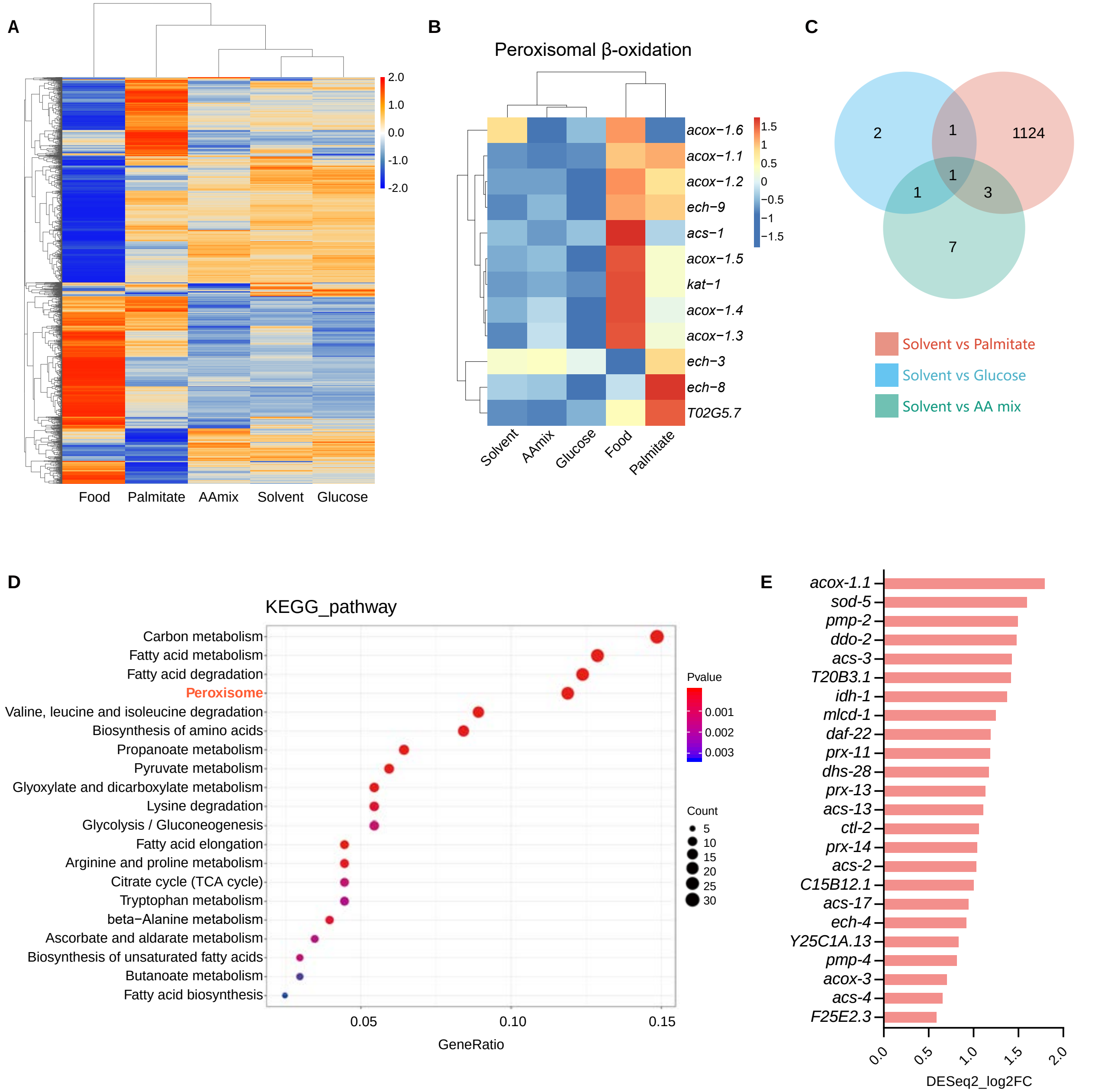


**Supplementary Figure 5.**

(A) A heatmap showing the gene expression of L1 animals under various nutrient conditions.

(B) A heatmap showing  the peroxisomal β-oxidation related genes of L1 animals under various nutrient conditions.

(C) A Venn diagram of up-regulated genes (compared to the solvent group) among three different expression gene sets (DEGs) shown by three colors. 1124 genes were up-regulated in the solvent (DMSO) vs palmitate group.

(D) A chart of the KEGG pathway analysis of the 1124 genes of the VENN diagram (C). Peroxisome-related genes are highly enriched.

(E) Expression change of genes in the peroxisome group (red in S5B) were listed.

.

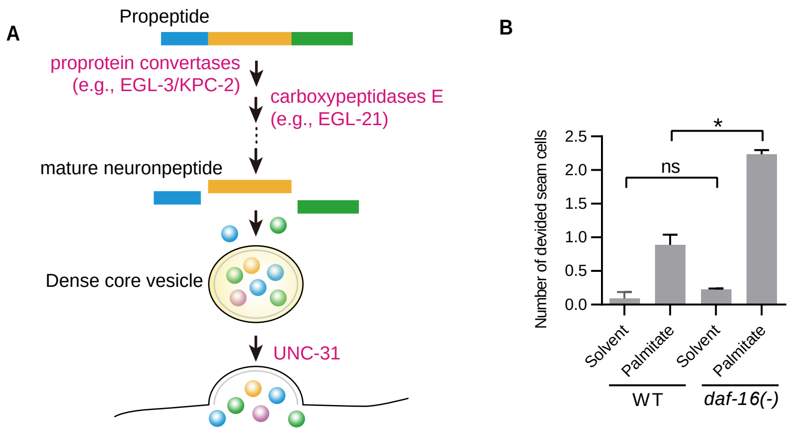


**Supplementary Figure 6.**

(A) A chart showing the neuropeptide processing and secretion pathway.

(B) A bar graph showing the average number of divided seam cells. A loss-of-function mutant of *daf-16* could not promote the division of seam cells by itself, but enhance the seam cell division robustly under palmitate supplementation.

.
