## Supplementary figures and images for "Free long chain fatty acid solitarily primes early postembryonic development in *Caenorhabditis elegans* under starvation"

### Supplementary tables

Table S1.

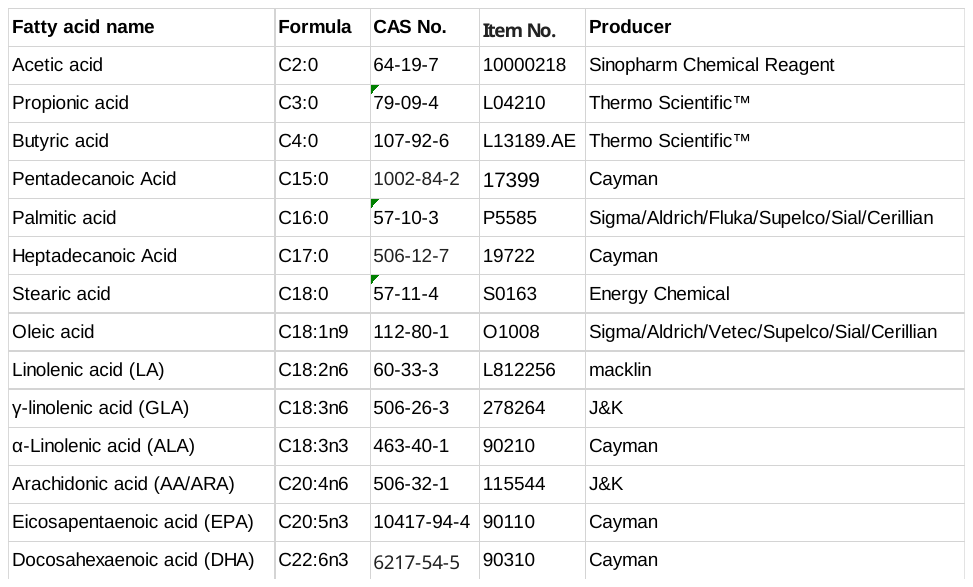

Table S2.

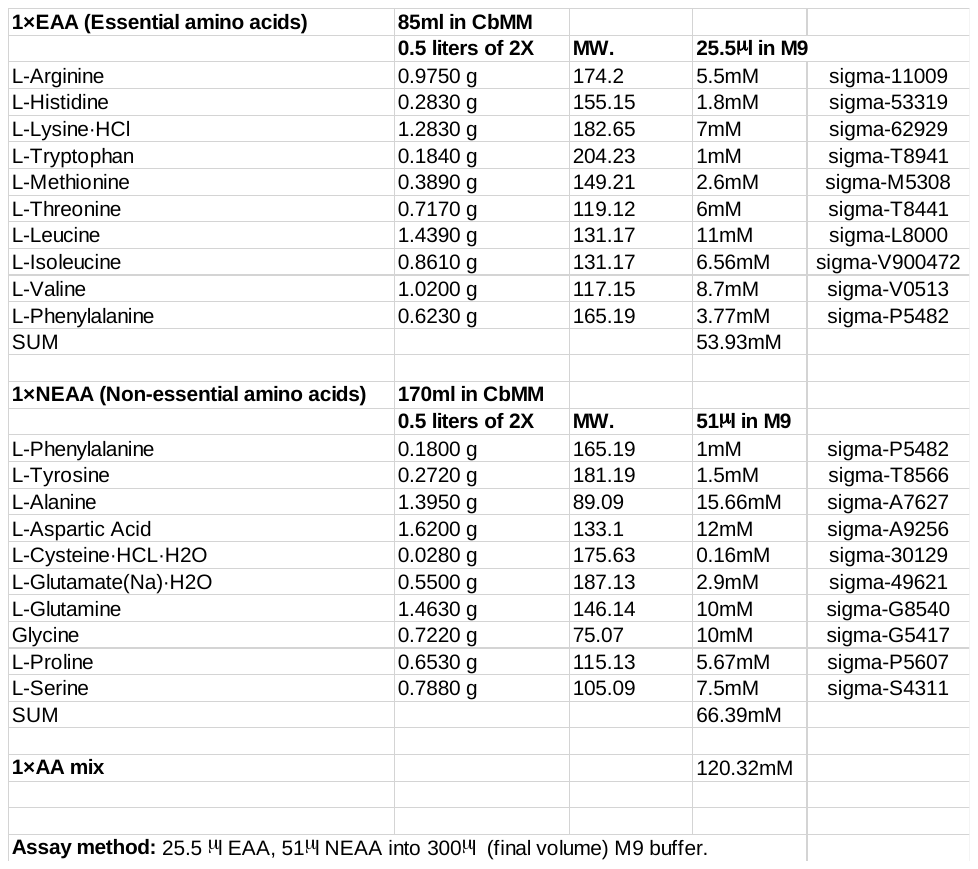
